# High-throughput affinity profiling of antibodies against soluble and native membrane targets by digital flow cytometry

**DOI:** 10.64898/2026.10.15.759441

**Authors:** Yuanhua Cheng, Sebastien Barcy, Bryant Fujimoto, Wyatt Nelson, Mengxia Zhao, Daniel T. Chiu

## Abstract

Binding affinity measurements are essential for optimizing and validating therapeutic and research antibodies, yet no high-throughput platform supports quantitative affinity measurements for both soluble and native membrane targets. Here we present a suite of digital flow cytometry (dFC) assays that determine equilibrium binding affinities by directly counting free and bound molecules in homogeneous mixtures without immobilization. The assays enable titration-free absolute *K*_D_ measurements for fluorescent binding partners, affinity measurements for unlabeled antibodies against membrane receptors on cell-derived extracellular vesicles (EVs), and multiplexed assays for rapid ranking or screening of antibody variants. We demonstrate applications by quantifying the effects of degree-of-labeling on the affinity of antibody-dye conjugates, and of drug conjugation on therapeutic antibody binding to receptors on human cancer EVs. By combining single-molecule quantitation, adaptable assay designs, and high-throughput analysis, dFC provides a versatile platform for therapeutic antibody optimization and research antibody validation.

## Introduction

Antibodies play central roles in therapeutics and in biomedical research. Advances in antibody technologies—including affinity maturation, humanization, bispecific antibodies, and antibody-drug conjugates (ADCs)^1,2^—have expanded the functionality of therapeutic antibodies,^3–5^ while conjugation of antibodies with fluorescent molecules, polymers, and nanoparticles has produced a vast catalog of analytical probes^6^ for biomedical research and clinical diagnostics. Because these antibody modifications can alter binding affinity^7^ and thereby affect therapeutic efficacy and analytical performance,^8^ quantitative characterization of binding affinity is essential for therapeutic antibody optimization and for validation of research antibodies and their conjugates. However, no existing affinity characterization method can provide high-throughput quantitative affinity measurements in both homogeneous solution and against membrane proteins in their native lipid environment. These limitations are significant because most therapeutic antibodies target cell surface proteins,^4^ as do many fluorescent antibody probes used in flow cytometry and tissue imaging, while incomplete characterization of antibodies and antibody-dye conjugates used in research has contributed to poor experimental reproducibility and a broader “antibody characterization crisis.” ^9–12^

Surface plasmon resonance (SPR)^13^ and biolayer interferometry (BLI)^14^ are among the most widely used technologies for antibody affinity characterization, providing label-free determination of association and dissociation rate constants (*k*_on_ and *k*_off_) from which the equilibrium dissociation constant (*K*_D_), the standard measure of binding affinity, is derived. However, SPR and BLI require the immobilization of one binding partner on a 2D solid-liquid interface, introducing artifacts such as steric hindrance, restricted molecular orientation, and mass transport limitations that can affect equilibrium measurements.^15–17^ Furthermore, these solid-phase methods cannot directly quantify antibody binding to membrane receptors in their native lipid environment.^18^ Alternate approaches that avoid immobilization include KinExA^19^ and conventional flow cytometry.^20,21^ KinExA is a solution-phase equilibrium method that can provide *K*_D_ values for ultra-high-affinity interactions, but cannot easily scale to high-throughput measurements. Conventional flow cytometry can evaluate antibody binding to receptors on intact cells with high throughput, but quantitative affinity measurements are limited by insufficient sensitivity and resolution, lack of standardized absolute quantitation, and high antibody consumption. Consequently, a critical gap remains for a technology capable of rapid quantitative affinity profiling of antibodies against both soluble targets and native membrane proteins.

To address these limitations, here we apply digital flow cytometry (dFC)—a single-molecule analytical platform featuring 100% single-fluorophore detection sensitivity and high particle and sample throughput^22^—to the quantitative affinity profiling of antibodies. Because dFC directly counts free and bound molecules simultaneously in homogeneous equilibrium mixtures, *K*_D_ can be calculated directly from the measured molecular populations without surface immobilization or wash steps, and in some cases without titration. Furthermore, because the assay does not require surface immobilization, it can be extended to membrane proteins presented in their native lipid bilayers on extracellular vesicles (EVs) or other lipid nanoparticles, allowing affinity measurements of unlabeled antibodies and antibody-drug conjugates (ADCs) via competition assays. Notably, EVs and biological nanoparticles represent a generalizable model system that can be engineered for the display of a broad range of functional membrane proteins.^23–26^ The rapid analysis, 12-channel multiplexing, and 96-sample format make dFC well suited for high-throughput screening and validation of antibody libraries. Additionally, dFC analysis does not require pure samples because binding events are registered based on the co-localization of two distinct fluorescence signals.

We previously developed dFC methods for measuring the absolute concentration and degree-of-labeling of antibody-dye conjugates^27^ and for multiparameter characterization of EVs.^28^ In this work, we extend dFC to the quantitative affinity profiling of antibodies. We demonstrate how digital quantitation of free and bound species enables a suite of complementary assay architectures encompassing fluorescent binding partners in homogeneous solution, unlabeled therapeutic antibodies binding membrane receptors presented on cell-derived EVs, and multiplexed assays for rapid ranking or screening of antibody variants. Together, these capabilities provide a versatile platform for therapeutic antibody optimization and research antibody validation.

## Results

### Overview of dFC affinity assay architectures

To enable quantitative affinity measurements across diverse antibody and target formats, we developed a suite of complementary dFC assay architectures that derive from a common underlying capability: digital quantitation of free and bound molecular species in homogeneous equilibrium mixtures (Figure 1):

(1) **Fluorescent, soluble binding partners:** Direct, titration-free determination of absolute *K*_D_ values by simultaneously counting free and bound molecules in a single equilibrium mixture.
(2) **Fluorescent antibody conjugates targeting membrane proteins:** Quantitative affinity measurements using equilibrium titration of fluorescent antibodies against membrane-dye-labeled, cell-derived EVs or other biological nanoparticles.
(3) **Fluorescent antibody variants:** Multiplexed competition assays for rapid determination of relative *K*_D_ for multiple fluorescent antibody variants (e.g., same antibody labeled with different fluorophores or different degrees of labeling).
(4) **Unlabeled antibodies and antibody**-**drug conjugates targeting membrane proteins:** Displacement assays involving titration of an unlabeled antibody or ADC and a fixed-concentration tracer (fluorescent antibody with known *K*_D_) against membrane dye-labeled, cell-derived EVs.
(5) **Unlabeled antibody and ADC libraries:** Simplified competition assays for rapidly measuring the relative affinities of unlabeled antibody or ADC variants; relative affinities can be converted to *K*_D_ values via a reference with known *K*_D_.

**Figure 1.**
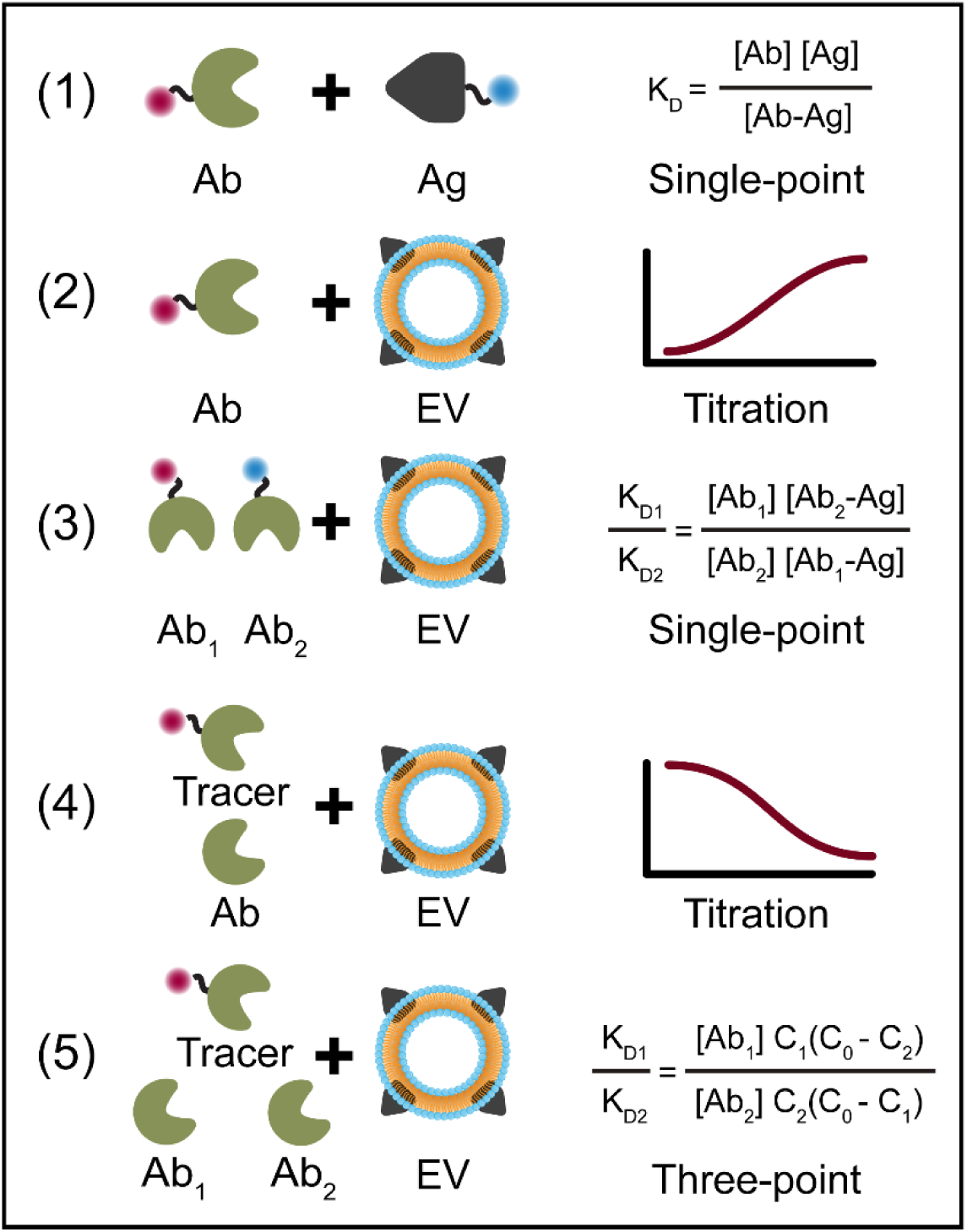
| Overview of dFC affinity assay architectures. Five complementary dFC assay formats for quantitative affinity measurements. (1) Single-point determination of absolute *K*_D_ for soluble fluorescent binding partners. (2) Equilibrium titration measurements of membrane-presented receptors using fluorescent antibody conjugates. (3) Multiplexed competition assay for relative affinity measurements of fluorescent antibody conjugates. (4) Competitive tracer assay for *K*_D_ measurements of unlabeled antibodies and antibody-drug conjugates (ADCs). (5) Three-point competition assay for rapid relative affinity measurements of unlabeled antibodies and ADCs.

The following sections describe the implementation and performance of each assay architecture.

### Titration-free absolute *K*_D_ measurements for fluorescent binding partners

The equilibrium dissociation constant, *K*_D_, is determined by the concentrations of free and bound species at equilibrium: *K*_D_ = [*A*][*B*]/[*AB*]. Conventional equilibrium methods generally measure only a single population—either bound complexes or free ligand—and therefore require titration series to estimate *K*_D_ from fitted binding curves. In contrast, in cases in which both binding partners are soluble and fluorescently labeled, dFC directly counts free binding partners and bound complexes simultaneously at the single-molecule level, enabling direct calculation of *K*_D_ from a single equilibrium sample without titration (Figure 2A).

**Figure 2.**
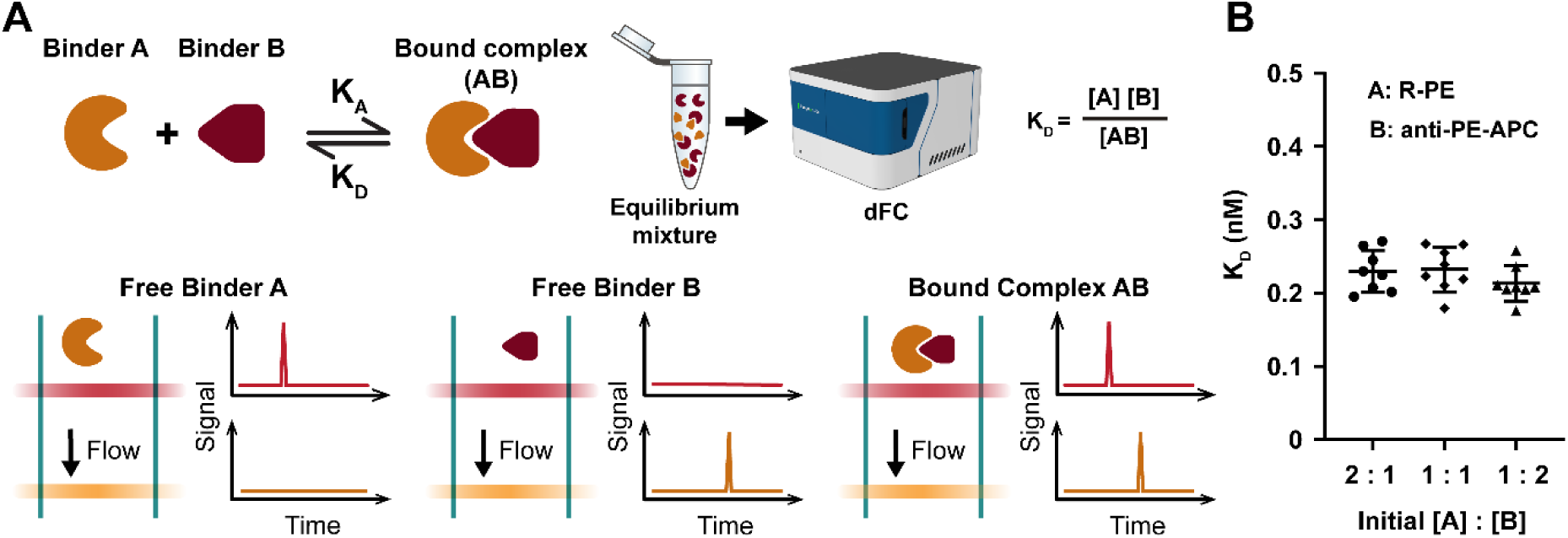
| Titration-free absolute *K*_D_ measurements for soluble fluorescent binding partners. (A) Schematic of the single-point affinity assay. Two fluorescently labeled binding partners are incubated to equilibrium and the resulting mixture is analyzed by dFC. Simultaneous digital counting of free binding partner A, free binding partner B, and the bound complex (AB) enables calculation of absolute *K*_D_ from a single sample. (B) Validation using phycoerythrin (PE) and anti-PE-APC. Similar *K*_D_ values were obtained across three initial PE:antibody molar ratios (2:1, 1:1, and 1:2), consistent with thermodynamic equilibrium.

This assay exploits two key capabilities of dFC. First, the ∼100% single-fluorophore detection sensitivity of dFC^22^ allows individual fluorescent molecules to be counted and converted to absolute molar concentrations through calibrated volumetric flow measurements. Second, when two fluorescently labeled binding partners form a complex, their coincident fluorescence signals are detected as single colocalized events, allowing free binder A, free binder B, and the bound complex (AB) to be quantified simultaneously (Figure 2A). From these three concentrations, *K*_D_ can be calculated from a single homogeneous equilibrium mixture without surface immobilization, wash steps, and/or curve fitting.

To validate this approach, we measured the interaction between an allophycocyanin (APC)-conjugated antibody and its target, the fluorescent protein phycoerythrin (PE). Direct counting of free antibody, free PE, and the bound antibody-PE complex yielded a *K*_D_ of 0.23 ± 0.03 nM (n = 8) from a single equilibrium mixture. Varying the initial PE:antibody ratio from 1:1 to 1:2 or 2:1 produced statistically indistinguishable *K*_D_ values (0.21 ± 0.02 nM and 0.23 ± 0.03 nM, both n = 8) (Figure 2B), indicating that the measured affinity was independent of the starting concentrations, as expected for an equilibrium measurement. Additionally, we examined the bound fractions of both PE and anti-PE in the three sets of equilibrium mixtures, and found the measured results are all consistent with the fractions predicted using the quadratic binding equation using the measured average *K*_D_ values (Figure S1).

This assay format is applicable to any binding interaction in which both binding partners are soluble and fluorescently labeled, and may be useful beyond antibodies for characterizing other proteins, peptides, nucleic acids, aptamers, glycans, and small-molecule interactions.

### *K*_D_ measurements for antibody-dye conjugates against membrane proteins

Many antibody-dye conjugates used in flow cytometry and histopathology target membrane receptors. To characterize their binding affinity while preserving the receptor’s native membrane environment, dFC employs equilibrium titration assays using cell-derived EVs (Figure 3A). Rather than labeling the receptor directly, EVs are stained with a lipid membrane dye (e.g., di-8-ANEPPS), enabling single-particle counting. Titration of a fluorescent antibody conjugate against a fixed concentration of labeled EVs allows colocalization analysis to identify antigen-positive vesicles and quantify the distribution of bound antibodies per EV (Figure 3A). The total number of bound antibodies is converted to the molar concentration of the bound state ([Ab-Ag]), and plotting this concentration against the total antibody concentration yields a saturation binding curve from which *K*_D_ is determined. Because these measurements are performed using membrane-presented receptors, the resulting affinity measurements are apparent *K*_D_ values which incorporate biologically relevant membrane effects such as receptor conformation, clustering, and avidity.

**Figure 3.**
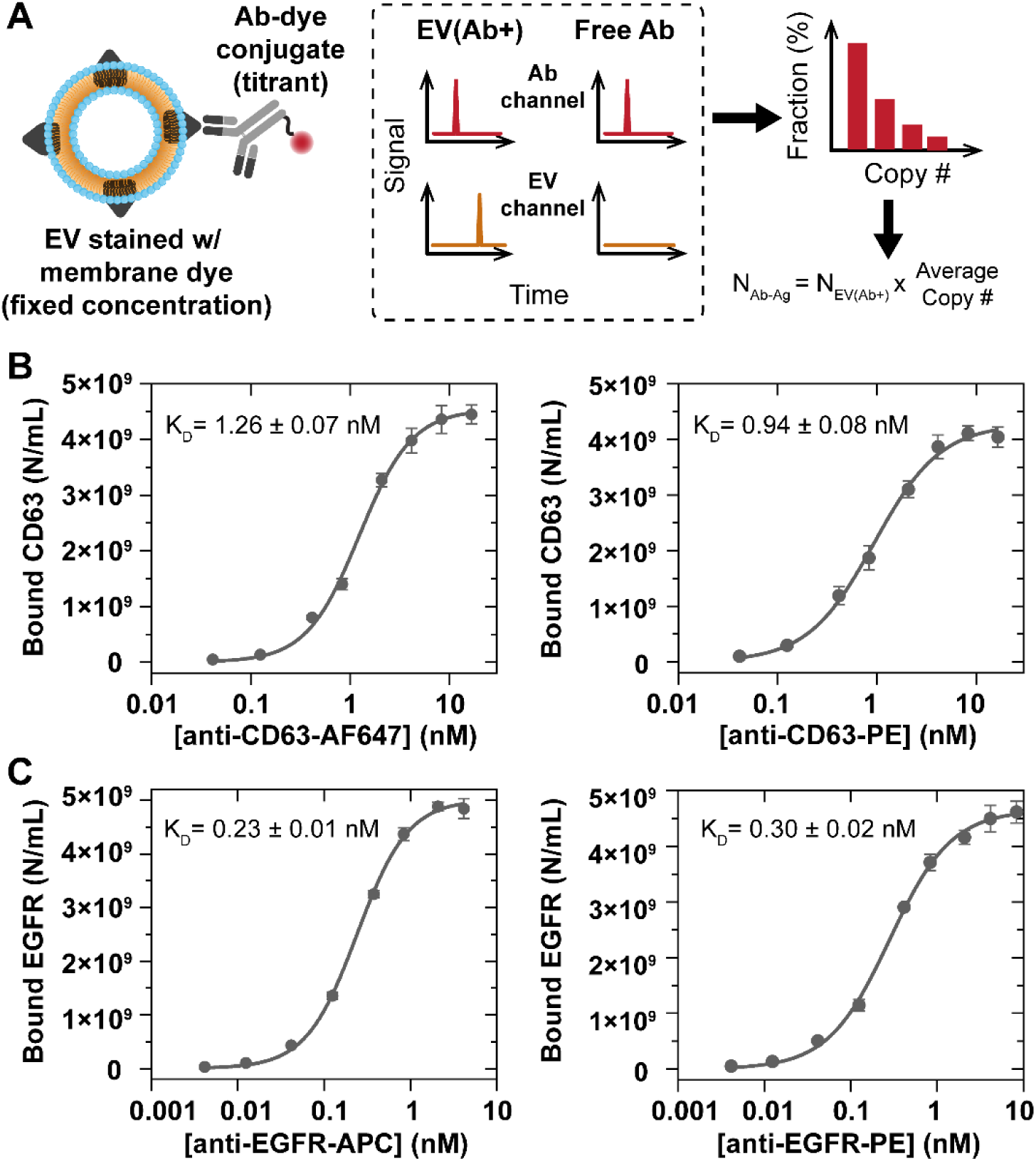
| *K*_D_ measurements for fluorescent antibody conjugates targeting membrane proteins. (A) Schematic of the equilibrium titration assay. Membrane-stained extracellular vesicles (EVs) with membrane receptors (antigens) are titrated with varying concentrations of an antibody-dye conjugate. Single-molecule colocalization identifies antigen-positive EVs and quantifies the number of bound antibodies per EV, allowing determination of the molar concentration of the bound state ([Ab-Ag]). (B) Titration curves and *K*_D_ values for two anti-CD63 antibody-dye conjugates, anti-CD63-AF647 (left) and anti-CD63-PE (right), binding to EVs. (C) Titration curves and *K*_D_ values for two anti-EGFR antibody-dye conjugates, anti-EGFR-APC (left) and anti-EGFR-PE (right), binding to EVs.

To demonstrate this approach, we used EVs derived from cultured cancer cells,^29^ and measured the binding affinities of antibody conjugates targeting CD63, a ubiquitous tetraspanin enriched on EV membranes. Titration of di-8-ANEPPS-stained EVs with anti-CD63-Alexa Fluor 647 (AF647) or anti-CD63-PE produced saturation binding curves yielding *K*_D_ values of 1.26 ± 0.07 nM and 0.94 ± 0.08 nM, respectively (n = 5) (Figure 3B). (The degree of labeling (DOL, also known as the fluorophore-to-protein ratio, F:P) of these conjugates, measured by dFC, was 2.36 ± 0.05 for CD63-AF647 and 1.15 ± 0.03 for CD63-PE. Below, we investigate the effect of DOL on affinity.)

We next examined antibody-dye conjugates targeting epidermal growth factor receptor (EGFR), a clinically important receptor displayed on the surface of EVs. Titration of di-8-ANEPPS-stained EVs with anti-EGFR (clone AY13) conjugated to APC (DOL, 1.09 ± 0.01) or PE (DOL, 1.22 ± 0.03) yielded *K*_D_ values of 0.23 ± 0.01 nM (APC) and 0.30 ± 0.02 nM (PE) (n = 5) (Figure 3C). These experiments demonstrate a quantitative equilibrium titration assay for determining *K*_D_ values of antibody-dye conjugates against membrane-presented receptors. The effects of fluorophore identity and DOL on the binding affinity of antibody-dye conjugates are further evaluated below using multiplexed competition assays.

### Rapid affinity profiling of antibody-dye conjugate variants

For rapid comparisons of antibody-dye conjugates consisting of the same antibody with different fluorophores or different degrees of labeling, a higher-throughput assay format is desirable. Therefore, we developed a multiplexed, titration-free competition assay that simultaneously determines the relative *K*_D_ values of multiple conjugates competing for a common target (Figure 4A). By directly quantifying the free and antigen-bound populations, relative affinities are determined in a single experiment without titration (Figure 4B).

**Figure 4.**
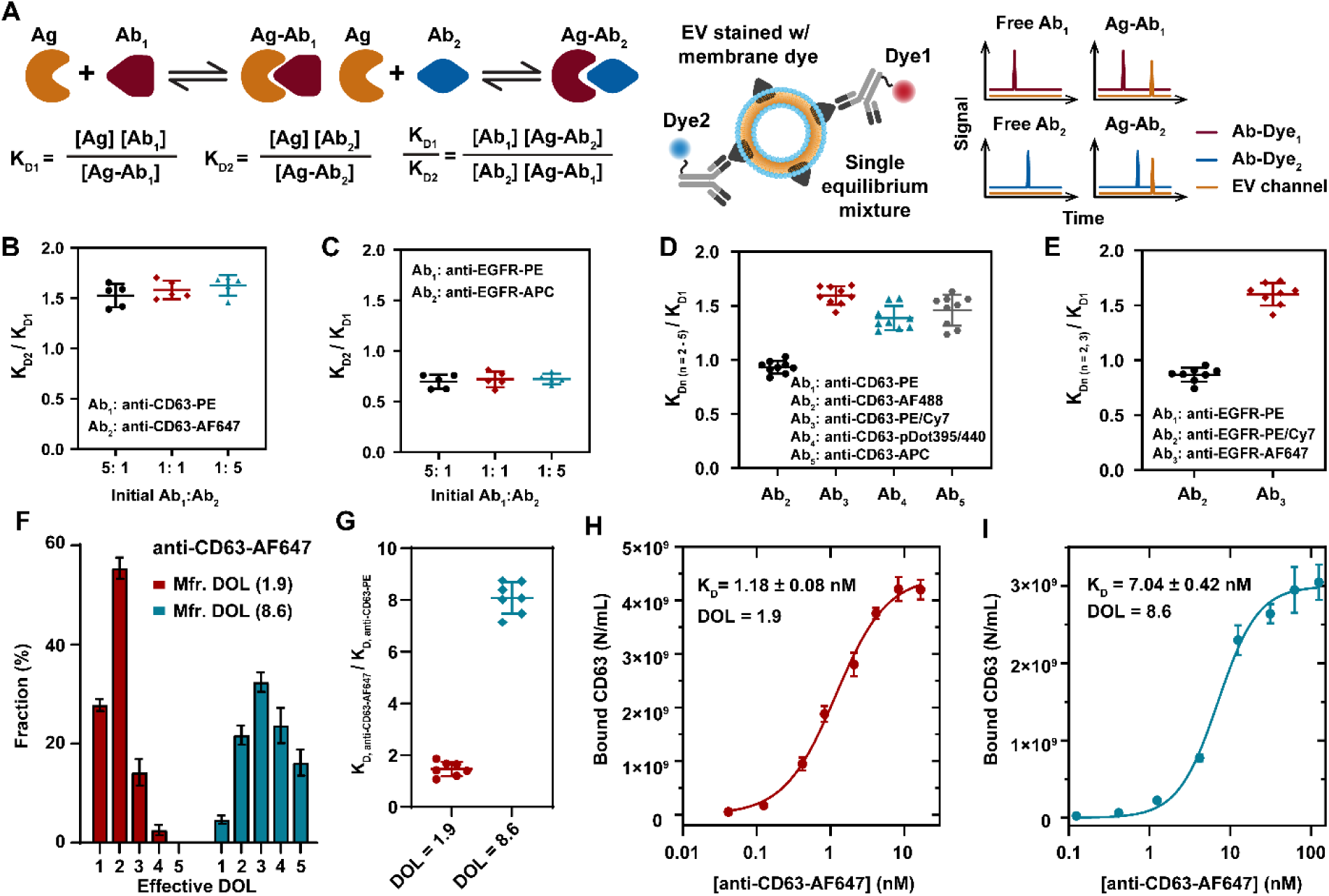
| Rapid affinity profiling of fluorescent antibody-dye conjugates. (A) Assay design. Left: Thermodynamic basis of the competition assay. Relative *K*_D_ values are determined from the free and antigen-bound populations of competing antibody conjugates. Right: dFC simultaneously quantifies the free and EV-bound populations of competing antibody conjugates in a single equilibrium mixture. (B, C) Relative *K*_D_ measurements of competing anti-CD63 (B) and anti-EGFR (C) conjugates across multiple initial concentration ratios. (D,E) Multiplexed relative affinity profiling of anti-CD63 (D) and anti-EGFR (E) conjugates labeled with different fluorophores. (F) Effective degree-of-labeling (DOL) distributions of two anti-CD63-AF647 conjugates. (G) Relative affinity measurements of the two anti-CD63-AF647 conjugates using anti-CD63-PE as the reference. (H, I) Equilibrium titration measurements of the low-(H) and high-DOL (I) conjugates, confirming the *K*_D_ values obtained by the competition assay.

To validate this approach, we performed a competition assay with the anti-CD63-PE and anti-CD63-AF647 conjugates described above, binding to CD63 on EVs. Simultaneous quantification of the free and EV-bound fractions of both conjugates yielded relative *K*_D_ values indicating that anti-CD63-PE binds approximately 1.5-fold more strongly than the AF647 conjugate (Figure 4B), in agreement with *K*_D_ values obtained by equilibrium titration (see above). Applying the assay to anti-EGFR-PE and anti-EGFR-APC likewise produced relative *K*_D_ values consistent with the titration-derived *K*_D_ values (Figure 4C and above). Varying the initial concentration ratio of the competing conjugates from 5:1 to 1:1 or 1:5 did not alter the measured relative affinities for either CD63 or EGFR, confirming that the assay reports equilibrium binding affinities. Together, these results establish a titration-free approach for rapid affinity profiling of antibody-dye conjugate variants.

We then applied the assay to compare the binding affinities of the same anti-CD63 antibody conjugated to five different fluorophores—the small-molecule dye AF488, the fluorescent proteins PE and APC, the protein tandem dye PE/Cy7, and the semiconducting polymer nanoparticle Pdot395/440— in a single experiment with EVs. Using the PE conjugate as the reference (relative *K*_D_ = 1), relative affinity measurements for the remaining conjugates were obtained with high reproducibility (Figure 4D). We similarly profiled anti-EGFR antibody (clone AY13) conjugated to PE, PE/Cy7, and AF647, with the PE conjugate as reference (Figure 4E). By using a reference conjugate with *K*_D_ measured by titration, relative affinity values can be quickly converted to *K*_D_ values for the entire panel.

This multiplexed competition assay can also be used to rapidly determine the effects of DOL on the binding of antibody conjugates. To demonstrate this capability, we analyzed anti-CD63-AF647 conjugates with DOL values of 1.9 ± 0.2 and 3.4 ± 0.4 (Figure 4F), measured by dFC using methods described previously.^27^ Applying the competition assay with anti-CD63-PE as the reference (relative *K*_D_ = 1), the conjugates exhibited relative *K*_D_ values of 1.41 ± 0.16 and 8.06 ± 0.62, respectively, indicating that the higher degree of labeling caused a >5-fold loss of antibody binding affinity (Figure 4G). Relative affinity values were converted to *K*_D_ values by referencing the known *K*_D_ of the PE conjugate, yielding *K*_D_ values of 1.32 ± 0.14 nM and 7.57 ± 0.58 nM, in close agreement with values obtained separately by titration (1.18 ± 0.08 nM and 7.04 ± 0.42 nM; Figure 4H-I). Together, these results establish a rapid, multiplexed approach for relative affinity profiling of antibody-dye conjugate variants, with *K*_D_ values recovered by reference to a single equilibrium titration measurement.

### *K*_D_ measurements for unlabeled therapeutic antibodies against membrane proteins

To enable quantitative affinity measurements of unlabeled therapeutic antibodies including antibody-drug conjugates (ADCs) which bind cell receptors, we developed a displacement titration assay in which an unlabeled antibody competes with a fixed-concentration fluorescent reference antibody (tracer) for binding to membrane receptors on EVs (Figure 5A). By quantifying the equilibrium redistribution of the tracer, dFC determines the binding affinity of the unlabeled antibody.

**Figure 5.**
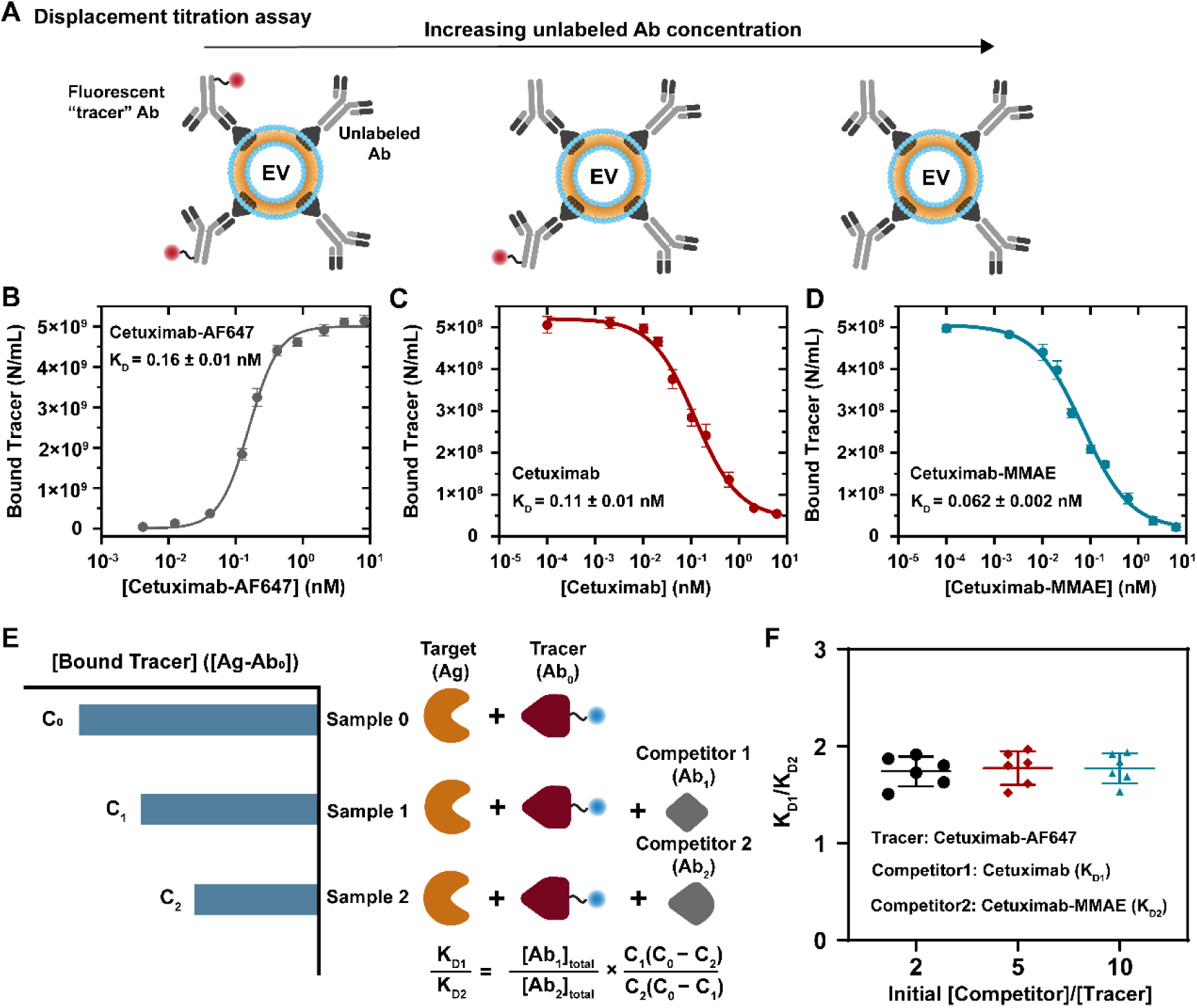
| *K*_D_ measurements for unlabeled therapeutic antibodies and ADCs. (A) Schematic of the displacement titration assay. An unlabeled antibody or antibody-drug conjugate (ADC) is titrated against a fixed concentration of fluorescent reference antibody (“tracer”) in equilibrium with EVs presenting the target membrane receptor. Redistribution of the tracer is used to determine the *K*_D_ of the unlabeled competitor. (B) Equilibrium titration (without competition; assay shown in Figure 3) to determine the *K*_D_ of the tracer, cetuximab-AF647. (C,D) Displacement titration of cetuximab (C) and cetuximab-MMAE (D) using cetuximab-AF647 as tracer. (E) Three-point competition assay for relative affinity measurements of unlabeled competitors. (F) Relative affinity measurements of cetuximab and cetuximab-MMAE across multiple competitor-to-tracer ratios.

To demonstrate this assay, we measured the binding affinity of cetuximab, an anti-EGFR therapeutic antibody approved for treatment of colorectal and head and neck cancers^30^ and its corresponding ADC, cetuximab-MMAE (monomethyl auristatin E), currently under preclinical investigation^31^ against EGFR presented on EVs. Cetuximab-AF647, with a *K*_D_ of 0.16 ± 0.01 nM (n = 5) determined by equilibrium titration (Figure 5B), was used as the fluorescent reference antibody (tracer) in competition assays with unlabeled cetuximab. Increasing concentrations of cetuximab produced progressive displacement of the tracer from EGFR-positive EVs (Figure 5C), yielding a *K*_D_ of 0.11 ± 0.01 nM (n = 5). This value agrees closely with the previously reported whole-cell ELISA affinity (*K*_D_ = 0.15 – 0.33 nM) ^32,33^ but is notably lower than affinities measured using SPR (*K*_D_ = 2-5 nM).^34,35^ Using the same assay, we measured the affinity of cetuximab conjugated to the small-molecule cytotoxin MMAE, obtaining a *K*_D_ of 0.062 ± 0.002 nM (n = 5) (Figure 5D), corresponding to an approximately 1.8-fold increase in binding affinity relative to the unconjugated antibody. These experiments demonstrate the competitive tracer assay as a quantitative approach for affinity measurements of unlabeled therapeutic antibodies and ADCs.

### Relative affinity measurements for rapid screening of unlabeled therapeutic antibodies

To enable affinity profiling of unlabeled antibodies without titration, we developed a simplified competition assay that determines the relative binding affinity of two unlabeled antibodies or ADCs competing for the same epitope using only three measurements (Figure 5E). A fluorescent tracer antibody is incubated with the target to establish the baseline concentration of bound tracer (*C*_0_). Two additional measurements are then performed using two competitor antibodies, yielding the residual tracer-bound concentrations *C*_1_ and *C*_2_. Under standard assay conditions in which the competitor concentration greatly exceeds the concentration of target binding sites, the relative affinity of the two competitors is calculated directly from these three measurements (Figure 5E and Supporting Information).

To validate this approach, we compared the relative binding affinities of cetuximab and cetuximab-MMAE, using cetuximab-AF647 as the fluorescent tracer (Figure 5F). Measured affinity ratios (*K*_D1_/*K*_D2_) were highly consistent across competitor-to-tracer molar ratios of 2:1, 5:1, and 10:1 (1.74 ± 0.15, 1.77 ± 0.17, and 1.77 ± 0.15; n = 6). These values closely matched the affinity ratio calculated from *K*_D_ values obtained separately by displacement titration (0.11 ± 0.01 nM / 0.062 ± 0.002 nM = 1.77 ± 0.17). Together, these experiments demonstrate the multiplexed competition assay for determining relative binding affinities of unlabeled antibodies without requiring a full titration. Utilized with the 96-flow-channel dFC chip, this assay will enable high-throughput library screening.

## Discussion

This work establishes digital flow cytometry (dFC) as a quantitative antibody affinity profiling platform for both soluble and membrane-presented targets. Building on the previously developed dFC system^22,27,28^ we developed a suite of equilibrium assay architectures for titration-free absolute *K*_D_ measurements, quantitative affinity measurements for native membrane-presented receptors, characterization of unlabeled therapeutic antibodies, and rapid multiplexed relative affinity measurements.

Current affinity characterization technologies each address only part of the antibody profiling landscape and have inherent limitations. SPR and BLI remain the predominant methods for antibody affinity characterization, providing association and dissociation rate constants which are used to calculate equilibrium dissociation constants, but are solid-phase methods that require surface immobilization. KinExA is a solution-phase equilibrium method that offers exceptional sensitivity and is compatible with ultra-high affinity binding^19^ but cannot be easily adapted to high throughput analysis. Conventional flow cytometry permits analysis of antibody binding to intact cells, but is often limited by insufficient sensitivity and resolution, inconsistent measurement results due to the lack of standardization and quality control, and high consumption of antibodies. Unlike these affinity measurement methods, which infer molecular populations from analog signals, dFC directly counts individual free and bound molecules, enabling absolute quantitation of the concentrations used to calculate *K*_D_. This capability is the foundation for adaptable assay designs including absolute *K*_D_ determination of fluorescent binding partners, characterization of unlabeled therapeutic antibodies binding to native membrane receptors, and rapid relative affinity profiling.

Another important capability of dFC is its ability to quantify antibody binding to membrane receptors in their native lipid environment. By using receptors presented on cell-derived EVs, these measurements capture biologically relevant determinants of binding, including membrane-dependent receptor conformation, post-translational modifications, receptor clustering, and avidity. Because the measurements are performed in a homogeneous, wash-free format, equilibrium is not perturbed during analysis. The close agreement between the cetuximab affinity measured by dFC and previously reported whole-cell ELISA measurements^32,33^ supports the biological relevance of this approach.

The assay architectures presented here provide practical capabilities for therapeutic antibody engineering and research reagent validation. The rapid sample analysis, multiplexed assays, and 96-sample microfluidic chip format provide high throughput. The competitive titration assays enable affinity measurements of unlabeled therapeutic antibodies binding membrane receptors, and the simplified three-measurement competition format enables efficient ranking or screening of libraries. These capabilities may allow rapid optimization of ADC design by quantifying the effects of drug-to-antibody ratio, linker chemistry, conjugation site, and payload on target binding affinity. The high-throughput profiling approach may also be useful for antibody-dye conjugate manufacturers to QC the vast libraries of conjugates required for quantitative flow cytometry and tissue imaging, and to help address the poor reproducibility of experimental results involving antibodies.^10,12^

Several limitations should be considered. The direct titration-free assay requires both binding partners to be fluorescently labeled and independently detectable. Membrane-based measurements report apparent *K*_D_ values that are specific to the biological context in which receptors are presented. Affinity measurements of unlabeled binders require an appropriately characterized fluorescent reference reagent. Future work will extend these assay architectures to additional classes of membrane particles, including lipid nanoparticles and viral vectors, and to biomolecular interactions beyond antibodies.

Together, these assays illustrate how digital quantification of free and bound molecular populations enables a broad range of affinity measurements encompassing both soluble and native membrane targets in a unified framework.

## Methods

### Digital flow cytometer (dFC)

The digital flow cytometer has been described and characterized previously.²² Four laser beams (405, 488, 561, and 637 nm; Coherent, CA) were spatially combined and shaped using a series of mirrors, cylindrical lenses, and convex lenses. The combined beams were focused onto the center of a microfluidic channel through a 40×, 0.95 NA air objective (Olympus, MA).

Fluorescence emission was collected through the same objective, separated from the excitation light by a multiband dichroic mirror (Chroma, VT), and spectrally divided using a series of dichroic mirrors and band-pass filters (Chroma, VT) before being focused onto 12 single-photon counting modules (SPCMs; Excelitas, Canada). Photon counts were acquired at 10,000 Hz using two digital counting cards (National Instruments, TX) and recorded with custom software developed in-house.

### Microfluidic chip fabrication

The microfluidic device was fabricated from polydimethylsiloxane (PDMS) using soft lithography as previously described.^22^ Microfluidic structures designed in AutoCAD (Autodesk, CA) were used to generate two high-resolution chrome photomasks (Front Range Photomask, NV). A two-layer silicon master was then fabricated. The first layer was produced by deep reactive ion etching to create features 2 µm in height, and the second layer was fabricated using SU-8 photoresist (MicroChem, MA) to generate features 30 µm in height. The dimensions of the detection channel were 2 × 2 × 125 µm (height × width × length). PDMS replicas molded from the silicon master were cured at 68 °C for 3 h and then bonded to a glass coverslip (Azer Scientific, PA).

### Direct measurement of absolute *K*_D_

This assay is for soluble fluorescent binding partners. To validate this approach, we used phycoerythrin (PE; Thermo Fisher Scientific, MA) and an allophycocyanin-conjugated anti-PE antibody (anti-PE-APC; BioLegend, CA) as a model system. After determining the stock concentrations by dFC, three equilibrium mixtures were prepared with PE:anti-PE-APC molar ratios of 1:1 (5 pM:5 pM), 2:1 (10 pM:5 pM), and 1:2 (5 pM:10 pM). All samples were prepared in PBS (pH 7.4; Gibco, CA) supplemented with 0.1% (w/v) BSA (Thermo Fisher Scientific, MA) and incubated at room temperature for 4 h. Finally, 10 µL of each equilibrium mixture was loaded into the microfluidic device for dFC analysis.

### Equilibrium titration assay for antibody-dye conjugates binding EVs

EVs were prepared as previously described.^28,29^ Antibody stock concentrations were measured by dFC before all titration experiments. For binding assays, 10 µL aliquots of EVs (∼2.5 × 10¹⁰ EVs/mL) suspended in PBS were incubated with target-specific antibody conjugates. Anti-CD63-AF647 and anti-CD63-PE (clone 5A6), together with a low-DOL anti-CD63-AF647 conjugate (DOL = 1.89), were titrated over eight final concentrations: 41.5 pM, 124.5 pM, 415 pM, 0.83 nM, 2.08 nM, 4.15 nM, 8.3 nM, and 16.6 nM. A high-DOL anti-CD63-AF647 conjugate (DOL = 8.57) was titrated over eight concentrations: 124.5 pM, 415 pM, 1.245 nM, 4.15 nM, 12.45 nM, 31.12 nM, 62.25 nM, and 124.5 nM. For anti-EGFR (clone AY13), the APC conjugate was titrated over eight concentrations: 4.15 pM, 12.45 pM, 41.5 pM, 124.5 pM, 415 pM, 0.83 nM, 2.08 nM, and 4.15 nM; the PE conjugate was titrated using an identical series with the addition of a ninth concentration at 8.3 nM. Cetuximab-AF647 (clone C225; Novus Biologicals, CO) was titrated over ten concentrations: 4.15 pM, 12.45 pM, 41.5 pM, 124.5 pM, 208 pM, 415 pM, 0.83 nM, 2.08 nM, 4.15 nM, and 8.3 nM.

Following antibody incubation, EV suspensions were stained with di-8-ANEPPS (100 nM final concentration; Thermo Fisher Scientific, MA) for 2 h at room temperature and purified by size-exclusion chromatography (Exo-Spin, 35-nm pore size; Cell Guidance Systems, MI). UltraPure BSA (Thermo Fisher Scientific, MA) was then added to a final concentration of 0.025% (w/v) to reduce nonspecific interactions. Finally, 10 µL of each sample was loaded into the microfluidic device for dFC analysis. For AF647– and APC-labeled samples, fluorescence was collected using the Ex405/Em600-50 and Ex488/Em600-50 channels for di-8-ANEPPS (to identify EVs by two-laser colocalization) and the Ex637/Em690-50 channel for AF647 or APC. For PE-labeled samples, fluorescence was collected using the Ex405/Em600-50 and Ex488/Em600-50 channels for di-8-ANEPPS and the Ex561/Em585-30 channel for PE.

### Multiplexed competition assay for fluorescent antibody conjugates

EVs were incubated with competing antibody pairs targeting either CD63 (anti-CD63-AF647 and anti-CD63-PE; clone 5A6) or EGFR (anti-EGFR-APC and anti-EGFR-PE; clone AY13). Aliquots of EVs (8 µL, ∼2.5 × 10¹⁰ EVs/mL) in PBS (pH 7.4) were incubated with the antibody pairs at three molar ratios. For CD63, the AF647 and PE conjugates were added at 0.167 nM/0.033 nM (5:1), 0.1 nM/0.1 nM (1:1), and 0.033 nM/0.167 nM (1:5). Similarly, anti-EGFR-APC and anti-EGFR-PE were incubated with EVs at 100 pM/20 pM (5:1), 60 pM/60 pM (1:1), and 20 pM/100 pM (1:5). Competitive profiling of two custom anti-CD63-AF647 conjugates with different DOL values was performed using anti-CD63-PE as the reference at 0.1 nM each. Following antibody incubation, EV suspensions were stained with di-8-ANEPPS (100 nM final concentration) for 2 h at room temperature. Fluorescence was acquired simultaneously using the Ex405/Em600-50, Ex488/Em600-50, Ex561/Em585-30, and Ex637/Em690-50 channels.

For multiplexed affinity profiling, EVs were incubated with two antibody panels: (1) anti-CD63 conjugated to AF488, PE, APC, PE/Cy7, and Pdot395/440; and (2) anti-EGFR conjugated to AF647, PE, or PE/Cy7. Aliquots of EVs (8 µL, ∼2.5 × 10¹⁰ EVs/mL) in PBS were incubated with each antibody conjugate at a final concentration of 50 pM together with di-8-ANEPPS (100 nM final concentration) for 2 h at room temperature. For the CD63 panel, fluorescence was acquired using the following seven channels: Ex405/Em434-22, Ex405/Em600-50, Ex488/Em520-30, Ex488/Em600-50, Ex561/Em585-30, Ex561/Em775-50, and Ex637/Em690-50. For the EGFR panel, fluorescence was acquired using the following five channels: Ex405/Em600-50, Ex488/Em600-50, Ex561/Em585-30, Ex561/Em775-50, and Ex637/Em690-50.

### Tracer displacement titration for unlabeled antibodies and antibody-drug conjugates

To measure the binding affinities of unconjugated cetuximab (MedChemExpress, NJ) and its corresponding antibody-drug conjugate, cetuximab-MMAE (MedChemExpress, NJ), 4 µL aliquots of EVs (∼2.5 × 10¹⁰ EVs/mL) in PBS were incubated with a fixed concentration of cetuximab-AF647 (20.5 pM) and varying concentrations of either cetuximab or cetuximab-MMAE. Both unlabeled competitors were titrated over an 11-point concentration series: 0, 0.1, 2.05, 10.25, 20.5, 41, 102.5, 205, and 615 pM, and 2.05 and 6.15 nM. Following antibody incubation, EV suspensions were stained with di-8-ANEPPS (100 nM final concentration) for 2 h at room temperature. Fluorescence was acquired using the Ex405/Em600-50, Ex488/Em600-50, and Ex637/Em690-50 channels.

### Three-point displacement assay

To compare the binding affinities of cetuximab and its corresponding antibody-drug conjugate, cetuximab-MMAE, three samples were prepared. Each sample contained EVs (∼2.5 × 10¹⁰ EVs/mL) and a fixed concentration of cetuximab-AF647 (20.5 pM; tracer) in PBS. The first sample served as the baseline tracer measurement and contained no competitor. The second and third samples contained 41 pM unlabeled cetuximab and 41 pM cetuximab-MMAE, respectively, corresponding to a competitor-to-tracer molar ratio of 2:1. Following antibody incubation, EV suspensions were stained with di-8-ANEPPS (100 nM final concentration) for 2 h at room temperature. Fluorescence was acquired using the Ex405/Em600-50, Ex488/Em600-50, and Ex637/Em690-50 channels. To validate the assay over a broader concentration range, additional measurements were performed at competitor concentrations of 102.5 pM (5:1) and 205 pM (10:1) for both cetuximab and cetuximab-MMAE.

### Data analysis and curve fitting

Raw photon counts were processed to identify and quantify individual fluorescent molecules passing through the laser using previously described methods.^22^ Baseline values were calculated from the mean background signal in each channel, with the corresponding noise approximated as the square root of the baseline value. Single fluorescent events were identified when photon bursts exceeded a signal-to-noise ratio (SNR) threshold which was optimized by comparing analyte signals with the corresponding blank control to maximize separation of true events from background noise.^22^ An autocorrelation function was applied to the di-8-ANEPPS signals acquired in the Ex405/Em600-50 and Ex488/Em600-50 channels to identify colocalized events corresponding to individual EVs. Based on the transit time of each EV between the 405– and 488-nm excitation lasers, signals detected in the remaining fluorescence channels were classified as EV-bound or unbound. Signals that did not colocalize with an EV were classified as unbound antibodies. The average number of antibodies bound per EV was determined by deconvolution using the unbound antibodies as an internal reference, as described previously.^27,28^ Briefly, the peak intensity distribution of unbound antibodies was modeled using a log-normal distribution, and the distribution of antibody copy numbers per EV was obtained by least-squares deconvolution of the bound and unbound fluorescence intensity distributions. The concentration of bound antibody was then calculated as: [Bound antibody] = [EV] × Fraction × Average copy number. Finally, *K*_D_ values from equilibrium titration experiments were determined by fitting the data to a quadratic binding model, and tracer displacement titration data for cetuximab and cetuximab-MMAE were fit using a sigmoidal dose-response model.

## Acknowledgements

We gratefully acknowledge support from the National Institutes of Health (R01DA057748 and R01HD113032) and the Stuart and Molly Sloan Precision Oncology Institute at the Fred Hutch Cancer Center through a Technology Dissemination Award. Part of this work was conducted at the Washington Nanofabrication Facility / Molecular Analysis Facility, a National Nanotechnology Coordinated Infrastructure (NNCI) site at the University of Washington with partial support from the National Science Foundation via awards NNCI-1542101 and NNCI-2025489.

