## Supporting Information for "High-throughput affinity profiling of antibodies against soluble and native membrane targets by digital flow cytometry"

### Table of Contents


Thermodynamic basis of the titration-free displacement assay **p. S3**

Experimental validation of effective DOL measurements **p. S7**

**Table S1.** Summary of dFC affinity profiling assays **p. S8**

**Table S2.** Summary of $K_D$ values for anti-CD63 antibody-dye conjugates **p. S9**

**Table S3.** Summary of $K_D$ values for anti-EGFR antibody-dye conjugates **p. S9**

**Table S4.** Summary of reagents **p. S10**

**Figure S1.** Experimental and calculated bound fractions in equilibrium mixtures **p. S11**

**References** **p. S12**

**Thermodynamic basis of the titration-free displacement assay**

**Assay design.** To determine the ratio of $K_D$ ($K_{D,1}/K_{D,2}$) between two non-fluorescent antibodies ($Ab_1$, and $Ab_2$) that compete for the same target antigen (Ag), we designed the following three samples using a fluorescent tracer antibody ($Ab_0$): **Sample 0** is the baseline measurement for the tracer ($Ab_0$) bound with the target antigen (Ag). It consists of one binding equilibrium, $Ab_0 + Ag \rightleftharpoons Ab_0\text{-}Ag$, with the input and equilibrium concentrations described below,

| | Tracer | Antigen | Bound Tracer |
|---|---|---|---|
| Input conc. | $[Ab_0]_{total}$ | $[Ag]_{total}$ | 0 |
| Equilibrium conc. | $[Ab_0]_0$ | $[Ag]_0$ | $[Ab_0\text{-}Ag]_0$ |

**Sample 1** is the competitive binding between the tracer ($Ab_0$) and the first unlabeled antibody ($Ab_1$) targeting the same antigen (Ag). It consists of two equilibria: $Ab_0$+Ag$\rightleftharpoons$$Ab_0$-Ag and $Ab_1$+Ag$\rightleftharpoons$$Ab_1$-Ag, with the input and equilibrium concentrations summarized below,

| | Tracer | Competitor | Antigen | Bound Tracer | Bound Competitor |
|---|---|---|---|---|---|
| Input conc. | $[Ab_0]_{total}$ | $[Ab_1]_{total}$ | $[Ag]_{total}$ | 0 | 0 |
| Equilibrium conc. | $[Ab_0]_1$ | $[Ab_1]_1$ | $[Ag]_1$ | $[Ab_0\text{-}Ag]_1$ | $[Ab_1\text{-}Ag]_1$ |

**Sample 2** is the competitive binding between the tracer ($Ab_0$) and the second unlabeled antibody ($Ab_2$) targeting the same antigen (Ag). It consists of two equilibria: $Ab_0$+Ag$\rightleftharpoons$$Ab_0$-Ag and $Ab_2$+Ag$\rightleftharpoons$$Ab_2$-Ag, with the input and equilibrium concentrations summarized below,

| | Tracer | Competitor | Antigen | Bound Tracer | Bound Competitor |
|---|---|---|---|---|---|
| Input conc. | $[Ab_0]_{total}$ | $[Ab_2]_{total}$ | $[Ag]_{total}$ | 0 | 0 |
| Equilibrium conc. | $[Ab_0]_2$ | $[Ab_2]_2$ | $[Ag]_2$ | $[Ab_0\text{-}Ag]_2$ | $[Ab_2\text{-}Ag]_2$ |

For all three samples, the input concentration of the antigen ($[Ag]_{total}$) is maintained at a fixed value, as is the input concentration of the tracer ($[Ab_0]_{total}$).

**Thermodynamic derivation**. In Sample 0, following the definition of dissociation constant ($K_D$), the binding affinity of the tracer ($Ab_0$) is,

$$K_{D,\,0}=\frac{[Ab_0]_0[Ag]_0}{[Ab_0\text{-}Ag]_0} \text{ (eq. 1)}$$

For antigen (Ag) in this equilibrium,

$$[Ag]_{total} = [Ag]_0 + [Ab_0\text{-}Ag]_0 \ (eq. 2)$$

Rearranging eq. 1 and 2, we obtained the classical Langmuir isotherm form of binding fractions:

$$[Ab_0\text{-}Ag]_0 = [Ag]_{total}\frac{\frac{[Ab_0]_0}{K_{D,\,0}}}{1+\frac{[Ab_0]_0}{K_{D,\,0}}} \text{ (eq. 3)}$$

Similarly, for Sample 1, we can obtain the following equations based on the definition of $K_D$ and the mass balance constraint,

$$K_{D,\,0} = \frac{[Ab_0]_1[Ag]_1}{[Ab_0\text{-}Ag]_1} \text{ (eq. 4)}$$

$$K_{D,\,1} = \frac{[Ab_1]_1[Ag]_1}{[Ab_1\text{-}Ag]_1} \text{ (eq. 5)}$$

$$[Ag]_{total} = [Ag]_1 + [Ab_0\text{-}Ag]_1 + [Ab_1\text{-}Ag]_1 \text{ (eq. 6)}$$

Combining these three equations, we get,

$$[Ag]_{total} = [Ag]_1 + \left(\frac{[Ab_0]_1[Ag]_1}{K_{D,\,0}}\right) + \left(\frac{[Ab_1]_1[Ag]_1}{K_{D,\,1}}\right) \text{ (eq. 7)}$$

Rearranging eq. 7, we obtain the following equation,

$$[Ag]_1 = \frac{[Ag]_{total}}{1+\frac{[Ab_0]_1}{K_{D,\,0}}+\frac{[Ab_1]_1}{K_{D,\,1}}} \text{ (eq. 8)}$$

The bound fraction of tracer in Sample 1 ($[Ab_0\text{-}Ag]_1$) would thus be,

$$[\mathrm{Ab_0\text{-}Ag}]_1 = [\mathrm{Ag}]_1 \frac{[\mathrm{Ab_0}]_1}{\mathrm{K_{D,\,0}}} = [\mathrm{Ag}]_{\mathrm{total}} \left( \frac{\frac{[\mathrm{Ab_0}]_1}{\mathrm{K_{D,\,0}}}}{1 + \frac{[\mathrm{Ab_0}]_1}{\mathrm{K_{D,\,0}}} + \frac{[\mathrm{Ab_1}]_1}{\mathrm{K_{D,\,1}}}} \right) \text{(eq. 9)}$$

Combining eq. 3 and eq. 9, we can obtain the change in the concentration of bound tracer between Sample 0 and 1,

$$[\mathrm{Ab_0\text{-}Ag}]_0 - [\mathrm{Ab_0\text{-}Ag}]_1 = [\mathrm{Ag}]_{total} \left( \frac{\frac{[Ab_0]_0}{K_{D,0}}}{1 + \frac{[Ab_0]_0}{K_{D,0}}} - \frac{\frac{[\mathrm{Ab_0}]_1}{\mathrm{K_{D,\,0}}}}{1 + \frac{[\mathrm{Ab_0}]_1}{\mathrm{K_{D,\,0}}} + \frac{[\mathrm{Ab_1}]_1}{\mathrm{K_{D,\,1}}}} \right)$$

$$= \frac{[Ag]_{total}}{K_{D,0}} \left( \frac{[Ab_0]_0 + \frac{[Ab_0]_0[\mathrm{Ab_1}]_1}{\mathrm{K_{D,\,1}}} - [Ab_0]_1}{\left(1 + \frac{[Ab_0]_0}{K_{D,0}}\right)\left(1 + \frac{[\mathrm{Ab_0}]_1}{\mathrm{K_{D,\,0}}} + \frac{[\mathrm{Ab_1}]_1}{\mathrm{K_{D,\,1}}}\right)} \right) (eq. 10)$$

Divide eq. 10 by eq. 9, we obtain,

$$\frac{[\mathrm{Ab_0\text{-}Ag}]_0 - [\mathrm{Ab_0\text{-}Ag}]_1}{[\mathrm{Ab_0\text{-}Ag}]_1} = \frac{[Ab_0]_0 + \frac{[Ab_0]_0[\mathrm{Ab_1}]_1}{\mathrm{K_{D,\,1}}} - [Ab_0]_1}{\left(1 + \frac{[Ab_0]_0}{K_{D,0}}\right)[Ab_0]_1} \quad (eq. 11)$$

Similarly, we can obtain the following equation for sample 2,

$$\frac{[\mathrm{Ab_0\text{-}Ag}]_0 - [\mathrm{Ab_0\text{-}Ag}]_2}{[\mathrm{Ab_0\text{-}Ag}]_2} = \frac{[Ab_0]_0 + \frac{[Ab_0]_0[\mathrm{Ab_2}]_2}{\mathrm{K_{D,\,1}}} - [Ab_0]_2}{\left(1 + \frac{[Ab_0]_0}{K_{D,0}}\right)[Ab_0]_2} \quad (eq. 12)$$

For these three samples,

$$[Ab_0]_0 = [Ab_0]_{total} - [\mathrm{Ag\text{-}Ab_0}]_0$$

$$[Ab_0]_1 = [Ab_0]_{total} - [\mathrm{Ag\text{-}Ab_0}]_1$$

$$[Ab_0]_2 = [Ab_0]_{total} - [\mathrm{Ag\text{-}Ab_0}]_2$$

Because $[Ab_0]_{total} \gg [Ag]_{total}$,

$$[Ab_0]_0 \approx [Ab_0]_1 \approx [Ab_0]_2 \approx [Ab_0]_{total}$$

With this approximation, Eq. 11 and 12 are thus simplified as follows,

$$\frac{[\mathrm{Ab_0\text{-}Ag}]_0 - [\mathrm{Ag\text{-}Ab_0}]_1}{[\mathrm{Ab_0\text{-}Ag}]_1} = \frac{\frac{[\mathrm{Ab_1}]_1}{\mathrm{K_{D,\,1}}}}{\left(1 + \frac{[Ab_0]_0}{K_{D,0}}\right)} \quad (eq. 13)$$

$$\frac{[\mathrm{Ab_0\text{-}Ag}]_0 - [\mathrm{Ag\text{-}Ab_0}]_2}{[\mathrm{Ab_0\text{-}Ag}]_2} = \frac{\frac{[\mathrm{Ab_2}]_2}{\mathrm{K_{D,\,2}}}}{\left(1 + \frac{[Ab_0]_0}{K_{D,0}}\right)} \quad (eq. 14)$$

Dividing eq. 13 by eq. 14, we can obtain,

$$\frac{\frac{[\mathrm{Ab_0\text{-}Ag}]_0 - [\mathrm{Ag\text{-}Ab_0}]_1}{[\mathrm{Ab_0\text{-}Ag}]_1}}{\frac{[\mathrm{Ab_0\text{-}Ag}]_0 - [\mathrm{Ag\text{-}Ab_0}]_2}{[\mathrm{Ab_0\text{-}Ag}]_2}} = \frac{\frac{[\mathrm{Ab_1}]_1}{\mathrm{K_{D,\,1}}}}{\frac{[\mathrm{Ab_2}]_2}{\mathrm{K_{D,\,2}}}} \quad (eq. 15)$$

Rearranging eq. 15, we thus obtain the ratio between $\mathrm{K_{D,\,1}}$ and $\mathrm{K_{D,\,2}}$,

$$\frac{\mathrm{K_{D,\,1}}}{\mathrm{K_{D,\,2}}} = \frac{[\mathrm{Ab_1}]_1}{[\mathrm{Ab_2}]_2} \frac{[\mathrm{Ab_0\text{-}Ag}]_1}{[\mathrm{Ab_0\text{-}Ag}]_2} \frac{([\mathrm{Ab_0\text{-}Ag}]_0 - [\mathrm{Ab_0\text{-}Ag}]_2)}{([\mathrm{Ab_0\text{-}Ag}]_0 - [\mathrm{Ab_0\text{-}Ag}]_1)} \quad (eq. 16)$$

In Samples 1 and 2, the initial concentrations of antibodies are the same and much higher than the EV concentration, so at equilibrium,

$$[Ab_1]_1 \approx [Ab_1]_{total}$$

$$[Ab_2]_2 \approx [Ab_2]_{total}$$

With these approximations,

$$\frac{\mathrm{K_{D,\,1}}}{\mathrm{K_{D,\,2}}} = \frac{[\mathrm{Ab_1}]_{\mathrm{total}}}{[\mathrm{Ab_2}]_{\mathrm{total}}} \frac{[\mathrm{Ab_0\text{-}Ag}]_1}{[\mathrm{Ab_0\text{-}Ag}]_2} \frac{([\mathrm{Ab_0\text{-}Ag}]_0 - [\mathrm{Ab_0\text{-}Ag}]_2)}{([\mathrm{Ab_0\text{-}Ag}]_0 - [\mathrm{Ab_0\text{-}Ag}]_1)} \quad (eq. 17)$$

Using eq. 17, we could calculate the relative binding affinities between $Ab_1$ and $Ab_2$ using results from three separate experiments.

**Experimental validation of effective DOL measurements**

The effective degree of labeling (DOL) values of anti-CD63-AF647, anti-CD63-PE, anti-EGFR-PE and anti-EGFR-APC conjugates were measured as described previously.[1] Briefly, the stock concentrations of antibody-dye conjugates and the corresponding free dyes (i.e. biotin-AF647, R-PE, and APC) were first determined by dFC.[2] These stock samples were then diluted to a final concentration of ~5 pM, using phosphate-buffered saline (PBS, pH 7.4, Gibco, CA) with 0.1% (w/v) Pluronic F-127 (Thermo Fisher Scientific, MA) and 0.05% (w/v) bovine serum albumin (BSA; Thermo Fisher Scientific, MA). After sample dilution, 10 μL of each sample was loaded into the inlet reservoirs on the microchip, and single-molecule counting data were collected with the vacuum at -0.5 psi. The peak intensity distribution for the free dyes was then analyzed using a log-normal fitting model. The distribution of copy number values was obtained from the deconvolution between the antibody-dye conjugates and the corresponding free dyes using a least-square method.

**Table S1. Summary of dFC affinity profiling assays**

| Assay Type | Sample Type | Labeling Requirement | Number of Measurements Needed | Output | Use Case |
|---|---|---|---|---|---|
| Direct $K_D$ measurement | Two fluorescent soluble binders | Both binders labeled | One | Absolute $K_D$ | Characterization of Ab-dye conjugates |
| Equilibrium titration | Fluorescent antibody conjugate + antigen on EVs | Antibody labeled | Full titration | Apparent $K_D$ | Characterization of Ab-dye conjugates binding membrane proteins |
| Equilibrium competition | Same antibody conjugated with different dyes + antigen on EVs | All competitors labeled | One | Relative $K_D$ | High-throughput screening or QC of Ab-dye conjugates |
| Displacement titration | Unlabeled antibody + dye conjugate + antigen on EVs | Tracer only | Full titration | Apparent $K_D$ | Characterize therapeutic Abs and ADCs against native membrane receptors |
| Three-point displacement | Unlabeled antibodies + dye conjugate + antigen on EVs | Tracer only | Three | Relative $K_D$ | High-throughput screening of libraries of therapeutic Abs and ADCs against native membrane receptors |

**Table S2. Summary of $K_D$ values for anti-CD63 antibody-dye conjugates**

| Reagent | $K_D$ by titration assay (nM) | $K_D$ by competition assay | |
|---|---|---|---|
| | | Relative $K_D$[#] | $K_D$ (nM) |
| anti-human CD63, PE | 0.94 ± 0.08 | 1 | 0.94 ± 0.08 |
| anti-human CD63, AF647 (catalog version) | 1.26 ± 0.07 | 1.58 ± 0.09 | 1.46 ± 0.08 |
| anti-human CD63, PECy7 | n.a. | 1.59 ± 0.09 | 1.49 ± 0.08 |
| anti-human CD63, AF488 | n.a. | 0.92 ± 0.07 | 0.86 ± 0.07 |
| anti-human CD63, Pdot395/440 | n.a. | 1.39 ± 0.11 | 1.32 ± 0.11 |
| anti-human CD63, APC | n.a. | 1.46 ± 0.13 | 1.37 ± 0.13 |
| anti-human CD63, AF647 (customized, F/P: 1.90) | 1.18 ± 0.08 | 1.41 ± 0.16 | 1.32 ± 0.14 |
| anti-human CD63, AF647 (customized, F/P: 8.57) | 7.04 ± 0.42 | 8.06 ± 0.62 | 7.57 ± 0.58 |

[#] Relative $K_D$ values are referenced to anti-human CD63, PE

**Table S3. Summary of $K_D$ values for anti-EGFR antibody-dye conjugates**

| Reagent | $K_D$ by titration assay (nM) | $K_D$ by competition assay | |
|---|---|---|---|
| | | Relative $K_D$[#] | $K_D$ (nM) |
| anti-human EGFR, PE | 0.30 ± 0.02 | 1 | 0.30 ± 0.02 |
| anti-human EGFR, APC | 0.23 ± 0.01 | 0.72 ± 0.08 | 0.22 ± 0.02 |
| anti-human EGFR, PECy7 | n.a. | 0.87 ± 0.07 | 0.26 ± 0.02 |
| anti-human EGFR, AF647 | n.a. | 1.60 ± 0.10 | 0.48 ± 0.03 |

[#] Relative $K_D$ values are referenced to anti-human EGFR, PE

**Table S4.** Summary of reagents

| Reagent | Supplier | Catalog No. | Lot |
|---|---|---|---|
| PE anti-human CD63 antibody, Mouse IgG1, κ | BioLegend | 353003 | B457715 |
| Alexa Fluor® 647 anti-human CD63 antibody, Mouse IgG1, κ | BioLegend | 353015 | B368377 |
| Alexa Fluor® 488 anti-human CD63 antibody, Mouse IgG1, κ | BioLegend | 3530037 | B315692 |
| APC anti-human CD63 antibody, Mouse IgG1, κ | BioLegend | 353008 | B471997 |
| PECy7 anti-human CD63 antibody, Mouse IgG1, κ | BioLegend | 353010 | B422098 |
| Alexa Fluor® 647 anti-human CD63 antibody, Mouse IgG1, κ (Manufacturer F/P: 8.57) | BioLegend | 900010458 | B429596 |
| Alexa Fluor® 647 anti-human CD63 antibody, Mouse IgG1, κ (Manufacturer F/P: 1.90) | BioLegend | 900010458 | B429595 |
| PE anti-human EGFR antibody, Mouse IgG1, κ | BioLegend | 352904 | B462324 |
| APC anti-human EGFR antibody, Mouse IgG1, κ | BioLegend | 352905 | B469657 |
| PECy7 anti-human EGFR antibody, Mouse IgG1, κ | BioLegend | 352909 | B393849 |
| Alexa Fluor® 647 anti-human EGFR antibody, Mouse IgG1, κ | BioLegend | 352918 | B478460 |
| EGFR Antibody (C225 (Cetuximab)) [Alexa Fluor® 647] | Novus Biologicals | NBP2-75903AF647 | D207961 |
| Cetuximab | MedChemExpress | HY-P9905 | 728400 |
| Cetuximab MMAE | MedChemExpress | HY-171135 | 1171226 |
| Alexa Fluor™ 647-Biotin Control Peptide | Invitrogen | PV6032 | 3249231A |
| R-phycoerythrin | Invitrogen | P801 | 2303163 |
| APC [Allophycocyanin] | AAT bioquest | 2554 | 11009244 |
| APC anti-phycoerythrin (PE) Antibody | BioLegend | 408107 | B391966 |
| UltraPure™ BSA (50 mg/mL) | Thermo Fisher Scientific | AM2618 | 01333228 |
| Pluronic™ F-127, 0.2 µm filtered (10% in water) | Thermo Fisher Scientific | P6866 | 2555600 |
| Di-8-ANEPPS | Thermo Fisher Scientific | D3167 | 2438355 |

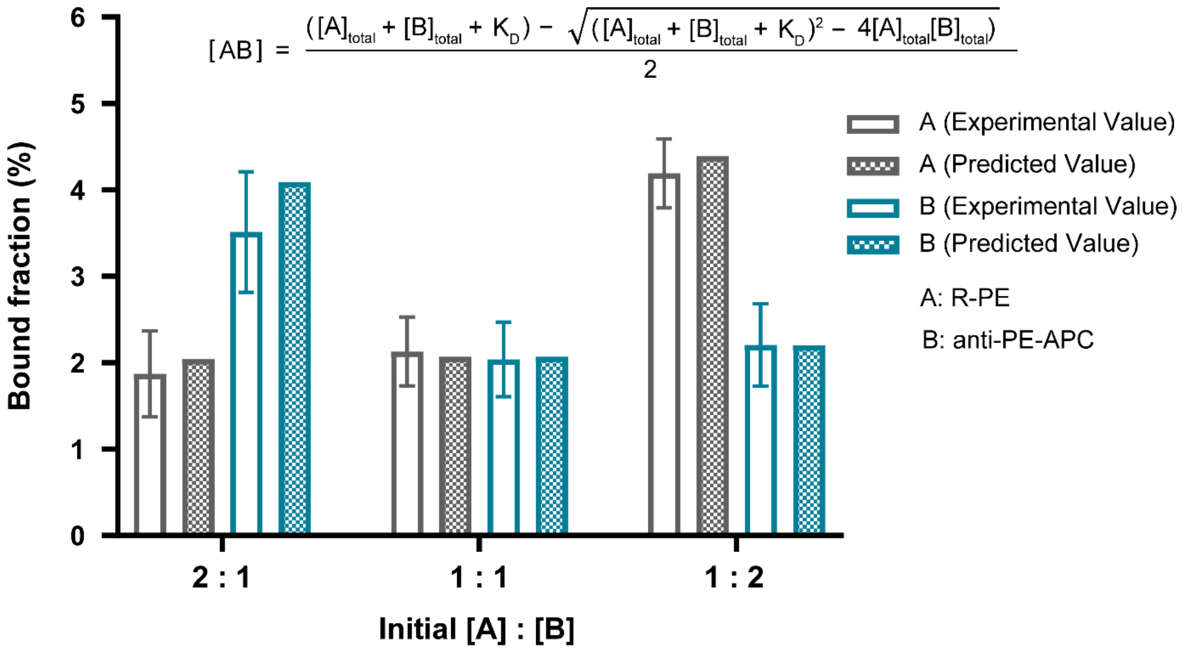


**Figure S1.** We measured the bound fractions (solid bars) of PE (grey) and anti-PE-APC (blue) in equilibrium mixtures with three sets of input molar concentrations: 1) $[PE]_{total}$ = 10 pM, $[anti\text{-}PE]_{total}$ = 5 pM, 2) $[PE]_{total}$ = 5 pM, $[anti\text{-}PE]_{total}$ = 5 pM, and 3) $[PE]_{total}$ = 5 pM, $[anti\text{-}PE]_{total}$ = 10 pM. Each bound fraction is defined as the ratio between the bound concentration and corresponding input concentration (e.g. Bound fraction of binder A = $[AB] / [A]_{total.}$ Error bars are standard deviations with 8 replicates in each set of samples. We also calculated the bound fractions at the equilibrium for the three sets of samples using the standard binding quadratic equation, with the corresponding input molar concentrations and the average $K_D$ values determined in Figure 2B. The measured bound fractions are consistent with the predicted values, suggesting our dFC method can accurately measure the molar composition of binding equilibrium.
